# solarius: an R interface to SOLAR for variance component analysis in pedigrees

**DOI:** 10.1101/035378

**Authors:** Andrey Ziyatdinov, Helena Brunel, Angel Martinez-Perez, Alfonso Buil, Alexandre Perera, Jose Manuel Soria

## Abstract

**Summary:** The open source environment R is one of the most widely used software for statistical computing. It provides a variety of applications including statistical genetics. Most of the powerful tools for quantitative genetic analyses are stand-alone free programs developed by researchers in academia. SOLAR is the standard software program to perform linkage and association mappings of the quantitative trait loci (QTLs) in pedigrees of arbitrary size and complexity. *solarius* allows the user to exploit the variance component methods implemented in SOLAR. It automates such routine operations as formatting pedigree and phenotype data. It also parses the model output and contains summary and plotting functions for exploration of the results. In addition, *solarius* enables parallel computing of the linkage and association analyses, that makes the calculation of genome-wide scans more efficient.

**Availability and implementation:** *solarius* is available on CRAN https://cran.r-project.org/package=solarius and on GitHub https://github.com/ugcd/solarius. See http://solar.txbiomedgenetics.org/ for more information about SOLAR.

## 1 Introduction

Variance component (VC) models or linear mixed models are powerful tools in genetic studies particularly of quantitative traits. These models are attractive because they account for the contribution of individual genetic locus, while efficiently including polygenic and other confounding effects shared among individuals. Implementation of the VC methods has traditionally been a computationally challenging task, and SOLAR is one of the first and well established VC tools that focuses on the analysis of quantitative trait loci (QTL) in extended pedigrees [1]. *solarius* delivers to the R user three main quantitative genetic models, polygenic, linkage and association, which are evaluated in SOLAR at the back-end and controlled via the user interface in R at the front-end.

**Table 1:**
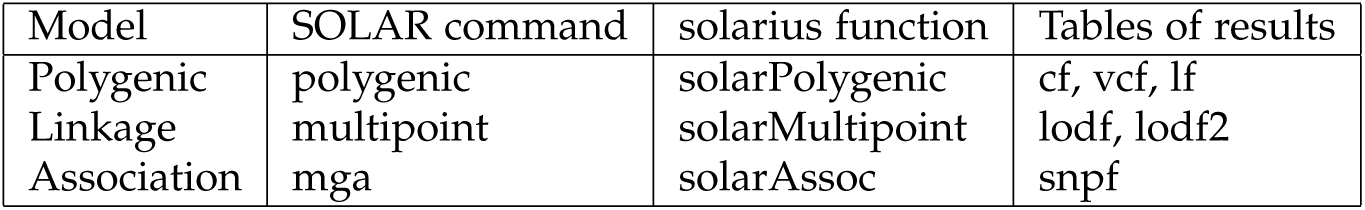
Implementation of the three main models in *solarius*. The high-level functions of the package (column 3) correspond to the low-level SOLAR commands (column 2). The results of an analysis are extracted from SOLAR output files and stored in slots of the returned objects in R (column 4). The main slots contain results for covariates (cf), variance components (vcf), likelihood statistics (lf), SNP associations (snpf), and logarithm of odds (LOD) scores (lodf and lodf2 for the first and the second passes, respectively).

The motivation to develop the *solarius* software came from the extensive experience of the group that studies the Genetic Analysis of Idiopathic Thrombophilia (GAIT) project [3, 4]. Both GAIT1 and GAIT2 projects include individuals from a Spanish population in extended pedigrees, an extensive list of clinical phenotypes related to the thrombotic disease and large arrays of both single nucleotide polymorphism (SNP) and microsatellite genetic markers. The first goal of the design of *solarius* was to provide an effortless data manipulation in a polygenic analysis needed to be explored for such large number of phenotypes. The second goal was to conduct the genome-wide scans for both linkage and association mappings in an efficient way by means of parallel computing.

## 2 Approach

### 2.1 Implementation

Our package automates the process of exporting data to the SOLAR-specific format, creating/cleaning directories, passing necessary configurations to SOLAR commands, and parsing SOLAR output files to extract the results of an analysis. The user works with top-level functions which correspond to low-level SOLAR commands, as summarized in Table 1. Each function performs the analysis with default SOLAR behavior, but the user can pass a specific configuration by two arguments, for example, `polygenic.settings and polygenic.options for solarPolygenic` function.

The back-compatibility with SOLAR is guaranteed by recording all commands passed to SOLAR and all output files from the polygenic analysis (solar slot of the returned objects). Additionally, the user can use dir argument to do all of the analysis in a specific directory instead of using a temporary directory (the default work flow).

The package has a number of benefits being a part of the R environment. The main functions returns output results as objects of S3 classes, for which print, summarize and plotting methods are defined. Pedigree relationships can be examined by `plotPed` and `plotKinship2` functions based on *kinship2* and *Matrix* R packages, respectively. The large tables of results from the association and linkage analyses are efficiently stored in *data.table* format from an R package of the same name. The results of the association analysis are explored with quantile-quantile (QQ) and Manhattan plots from *qqman* R package. In addition, *rsnps* R package can retrieve the information about the SNPs by sending queries to the NCBI database.

Implementation of parallel calculations is straightforward, since the association and linkage analyses are implicitly parallel problems, and the R environment offers a number of packages with parallel interfaces *(parallel, iterators* and *doParallel* packages). The user needs to introduce the only parameter cores (the number of cores) to configure parallel computing.

### 2.2 Reproducible examples

*solarius* contains simulated data sets of relatively small sample size, which can be used for reproducible demonstrations of how the main functions of the package work. For instance, simulated data adapted from *multic* R package [2] include 1200 individuals in 200 families, two traits trait1 and trait2 and two covariates age and sex, 6 identity by descent (IBD) multipoint matrices for Chromosomes 5. A smaller subset of 174 individuals in 29 families is available as dat30 data set.

~~~
data(dat30)
~~~

Call for the polygenic model with the formula interface is similar to that of the standard linear regression function lm.

~~~
modl <-solarPolygenic(trait1^~^1, dat30)
~~~

Either print or summary method applied to mod1 object shows that estimation of the heritability is 0.83 ± 0.10 with p-value 1.20 × 10^-10^. Both covariates appeared to be not statistically significant if the function is called again with `covtest` argument equal to `TRUE`.

The bivariate polygenic model partitions phenotypic variance observed in two traits `trait1` and `trait2` into genetic and environmental components. The model also introduces correlation coefficients per component between traits.

~~~
mod2 <-solarPolygenic(trait1+trait2^~^1, dat30, polygenic.options = ’-testrhoe -testrhog’)
~~~

To test the statistical significance of the correlation coefficients, `polygenic.options` argument is used. The genetic correlation between two traits is 0.97 ± 0.04 with p-value 2.25 × 10^-9^, and the environmental correlation is 0.41 ± 0.20 with p-value 0.14.

The linkage analysis of trait1 on Chromosome 5 gives the highest LOD score 3.56 at 3 cM position.

~~~
mibddir <-system.file(’extdata’, ’solarOutput’, ’solarMibdsCsv’, package = ’solarius’)
link <- solarMultipoint(trait1^~^1, dat30, mibddir = mibddir, chr = 5)
~~~

A hundred of synthetic SNPs were randomly generated for `dat30` data set. Two tables `genocovdat30` and `mapdat30` contain genotype data and annotation information, respectively. The only SNP `snp_86` shows a statistically significant association (p-value 9.10 × 10^-5^) after calling `solarAssoc` function to compute the model in parallel on two cores.

~~~
assoc <- solarAssoc(trait1 ^~^ 1, dat30, cores = 2, snpcovdata = genocovdat30, snpmap = mapdat30)
~~~

To explore the association results, functions `plotQQ` for `QQ-plot`, `plotManh` for Manhattan plot and annotate for SNP annotation are provided. More examples on polygenic, linkage and association analyses are available in the tutorial on-line http://ugcd.github.io/solarius/vignettes/tutorial.html.

## 3 Acknowledgment

The authors thank Professor W.H. Stone for revising the manuscript.

## Funding

This research was funded by the TEC2013-44666-R grant. This work was partially funded by the 2014SGR-2016 consolidated research group of the Generalitat de Catalunya, Spain. CIBER-BBN is an initiative of the Spanish ISCIII. This research was supported partially by grants PI-11/0184, PI-14/0582 and UIN2013-50833 from the Instituto Carlos III (Fondo de Investigación Sanitaria - FIS).

## Conflict of Interest

None declared.

## References

[1] L Almasy and J Blangero. Multipoint quantitative-trait linkage analysis in general pedigrees. American journal of human genetics, 62(5):1198–211, May 1998.

[2] Mariza De Andrade, Elizabeth J Atkinson, Christopher I Amos, and Jianfang Chen. Estimating Genetic Components of Variance for Quantitative Traits in Family Studies using the MULTIC routines. Technical report, Department of Epidemiology, U.T. MD Anderson Cancer Center Houston, Texas, 2006.

[3] J Blangero, J T Williams, and L Almasy. Novel family-based approaches to genetic risk in thrombosis. Journal of thrombosis and haemostasis : JTH, 1(7):1391–7, July 2003.

[4] Luis Vila, Angel Martinez-Perez, Mercedes Camacho, Alfonso Buil, Sonia Alcolea, Nuria Pujol-Moix, Marta Soler, Rosa Antón, Juan-Carlos Souto, Jordi Fontcuberta, and José-Manuel Soria. Heritability of thromboxane A2 and prostaglandin E2 biosynthetic machinery in a Spanish population. Arteriosclerosis, thrombosis, and vascular biology, 30(1):128–34, January 2010.

